# Synapse-related protein alterations and estradiol deficiency associate with early Parkinsonism in female A53T-α-synuclein transgenic mice fed on a high-fat diet

**DOI:** 10.1101/2025.10.22.678296

**Authors:** Swati Aggarwal, Ankit Biswas, Naman Kharbanda, Nitu Singh, Krishna Singh Bisht, Chandrayee Dey, Saloni Khatri, Arundhati Tiwari, Tushar Kanti Maiti

**Affiliations:** Laboratory of Functional Proteomics, Regional Centre for Biotechnology, NCR Biotech Science Cluster, Faridabad

**Keywords:** Parkinson’s disease, Sex differences, Proteomics, Estradiol, Neurodegeneration

## Abstract

Substantial evidence highlights the detrimental impact of a fat-rich diet on cognitive and emotional behaviour. Epidemiological studies have linked the consumption of saturated fat with an increased risk of Parkinson’s disease (PD), whereas a low-fat or ketogenic diet is reported to improve both motor and non-motor symptoms. Several animal model studies further support these associations. However, the impact of a high-fat diet (HFD) on sex-specific behavioural alterations and the underlying molecular mechanism in PD remains poorly studied. In the present study, we investigated the impact of HFD on PD progression in a sex-specific manner using the A53T transgenic mouse model of PD. Behavioural and pathophysiological analyses revealed a faster onset and progression of PD-like phenotype in female mice exposed to HFD compared with the male mice. Proteomics profiling of brain tissues demonstrated positive enrichment of immune system-related pathways in males, while females exhibited considerable downregulation of synapse-associated pathways under HFD conditions. The reduced estradiol level was identified as a potential factor contributing to synaptic dysfunction and the subsequent early onset of PD in female mice. These findings provide novel insights into the sex-specific consequences of HFD on PD pathogenesis and highlight the role of estrogen-linked synaptic vulnerability in mediating diet-induced PD onset.

## Introduction

Parkinson’s Disease (PD) is the second most common neurodegenerative disorder worldwide. According to the World Health Organisation (WHO), the prevalence of PD has doubled over the past 25 years, with an estimated 8.5 million individuals affected globally in 2019. The cardinal motor symptoms of PD include tremors, muscular rigidity, bradykinesia, and postural instability. In addition to these hallmark features, numerous non-motor symptoms, such as cognitive impairment, depression, anxiety, fatigue, gastrointestinal disturbances, and loss of smell or vision, frequently precede the motor impairments.^1^ The etiology of PD differs markedly between men and women, with the former exhibiting approximately 1.5 times higher prevalence. The major contributors to these sex-related differences include lifestyle, ageing, gene regulation, and gonadal hormones^2^. Estradiol, a major female sex hormone, has long been recognised for its neuroprotective effects. Notably, post-menopausal estrogen therapy has been associated with a reduced risk of developing PD^3–5^. Multiple estradiol-mediated neuroprotective pathways have been identified, including MAPK/ERK, PI3K/Akt, Wnt, RhoA/ROCK, mTORC1, and AMPK. These pathways are implicated in synaptogenesis, enhanced neuronal plasticity, inhibition of excitotoxicity and apoptosis, mitochondrial protection, and neurogenesis^6,7^.

In recent years, the influence of lifestyle factors and metabolic comorbidities such as diabetes on neurological disorders has gained significant attention. The detrimental effects of fat accumulation are well-documented in multiple psychiatric and neurological conditions, including depression, intellectual disability, multiple sclerosis, Alzheimer’s disease, and PD^8–11^. Emerging evidence indicates that early PD progression is linked with high-fat diet (HFD)-induced diabetes, primarily through the acceleration of pathological microvascular alterations^12^. Animal-based studies suggest that enhanced oxidative stress, neuroinflammation, and suppression of peroxisome proliferator-activated receptors (PPARS) are potential mechanisms underlying this association^13–16^.

Despite growing evidence, most of the animal studies in PD have been conducted exclusively in male mice. Given the inherent differences in body composition, the distinct impact of body mass index (BMI) on cognitive decline, antioxidant effects, and the presence of sex-specific hormones, it is critical to study the effect of HFD in both sexes.^17,18^ A recent study in male and female 3xTg-Alzheimer’s disease (AD) mice revealed marked sex-specific metabolic response, wherein male mice showed energy deficit and hypothalamic inflammation under a control diet, while females exhibited energy surplus. HFD feeding worsened the systemic inflammation in males but caused pronounced metabolic dysfunction in female mice, highlighting the differential impact of HFD in AD^19^. Although the underlying mechanisms remain unclear, similar sex-dependent effects have been observed in other models. In mThy1-hSNCA line 61 transgenic mice, HFD feeding led to metabolic dysregulation but conferred relative resistance to obesity, resulting in a lean phenotype in both sexes^20^. Together, these findings highlight critical knowledge gaps in the understanding of how HFD influences PD progression, particularly in a sex-specific context.

In this regard, we examined the sex-specific impact of HFD on the progression of PD using the A53T α-synuclein (αSyn) transgenic mouse model. The experimental design ensured that a group exhibiting an early PD phenotype was terminated alongside their age-matched groups for subsequent molecular characterisation. Notably, female mice displayed an early phenotype of PD after eight months of HFD feeding, initiated at 3.5 months of age, whereas male mice did not show such a phenotype at this time. Proteomic analysis of brain tissues further revealed distinct sex-dependent molecular signatures; pathways related to immune regulation and metabolism were predominantly enriched in males, while those associated with synaptic function were downregulated in females. Furthermore, a significant reduction in the level of estradiol suggests that disruption of estradiol-mediated regulation of synaptic proteins may contribute to early disease progression in females under HFD conditions ^21^

## Results

### Study Design

In this study, we used B6C3-Tg (prnp-SNCA *A53T)83vle/J transgenic mice, produced by the inbreeding of pairs obtained from Jackson Laboratory (USA). The Institutional Animal Ethics Committee of the Regional Centre for Biotechnology, Faridabad, approved all the experimental procedures. Genotyping was performed using the Jackson Laboratory protocol using qRT-PCR, and animals were categorised as homozygous, hemizygous, and heterozygous. Hemizygous mice were used as the PD group, while heterozygous mice served as the non-PD (NPD) group. Both males and females from each group were included in the study. After a two-week acclimatisation period, the 3.5-month-old mice were randomly assigned to either an HFD ( 60% fat, Research diet, D12492) or a control diet (10% fat, Research diet, D12450B). This resulted in eight experimental groups as follows: Non-PD control male (NPD-CM), Non-PD high-fat diet male (NPD-HM), PD control diet male (PD-CM ), PD high fat diet male (PD-HM ), Non-PD control diet female (NPD-CF ), Non-PD high fat diet female (NPD-HF), PD control diet female (PD-CF ) (Fig. 1). HFD feeding was continued until clear behavioural alterations indicative of PD onset were observed in any of the groups, at this point all age-matched groups were terminated for molecular and histological analyses.

**Fig. 1.**
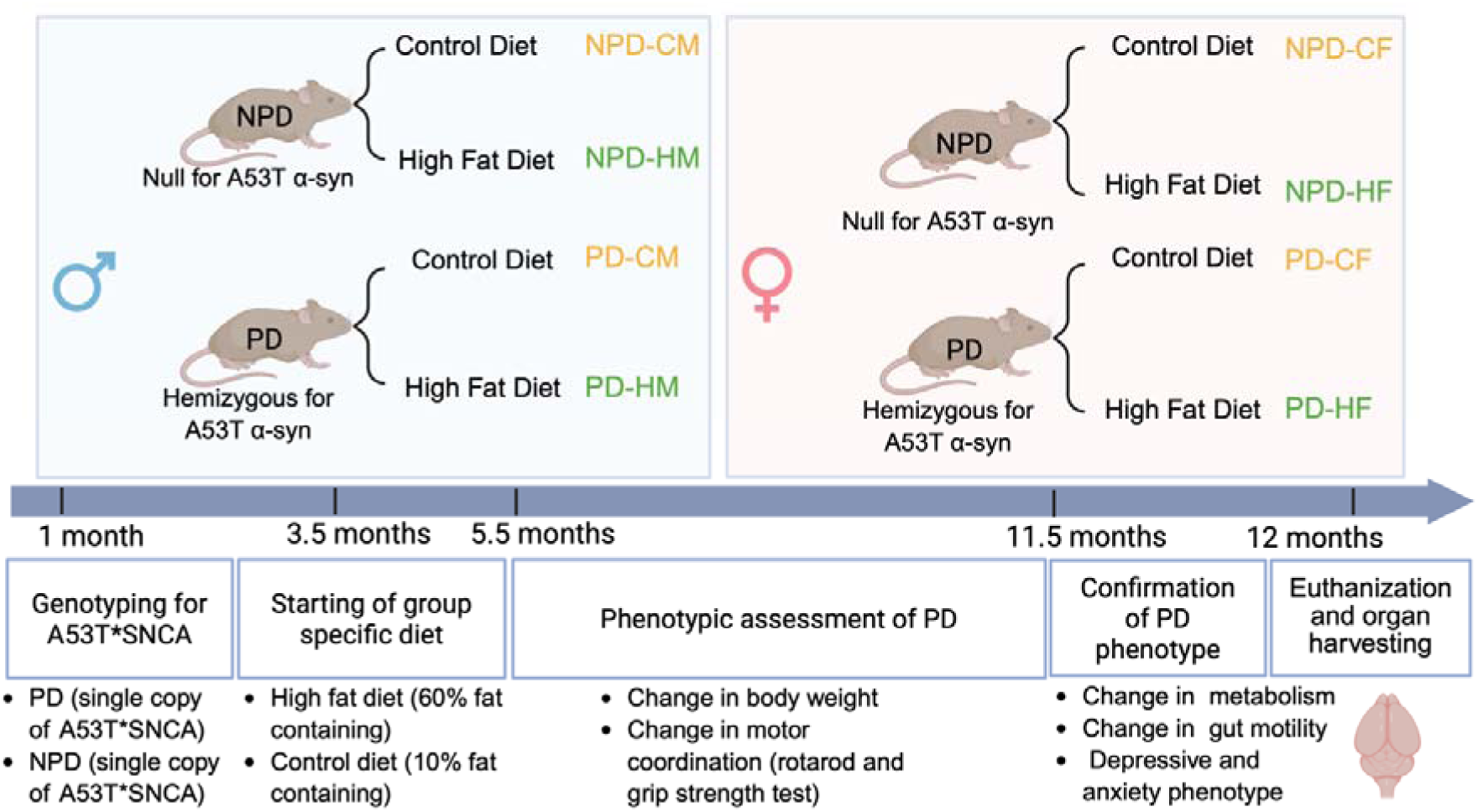
Development of a high-fat diet-induced Parkinsonism mouse model. A schematic representation of model development, along with a timeline indicating that mouse feeding was started from the age of 3.5 months, and phenotypic assessment of PD (rotarod and grip strength) reading was started from the age of 5.5 months and continued till the changes were not evident in any of the groups, that is, 11.5 months. While confirmatory phenotypic tests were conducted one week prior to the euthanisation (12M). The mouse brain and myenteric plexus were extracted and further used for molecular studies. This figure also indicates the grouping pattern based on biological sex, genotype, and feed administered, used in the study. Based on biological sex, there were 2 groups: male and female, which were further segregated based on the absence or presence of A53T mutant form of SNCA in their genotype, namely PD (Presence of a single copy of A53T SNCA, making mice prone to PD) and NPD (absence of A53T SNCA acting as Non-prone for PD). These were further sub-grouped based on the diet given: High-fat diet (60% fat) and control diet (10% fat), ultimately leading to 8 groups, which are nomenclature genotype (PD/NPD)-diet(C/H), biological sex(M/F).

### A high-fat diet alters metabolic homeostasis in mice

To assess the impact of diet on metabolic homeostasis, we monitored body weight changes throughout the study and measured respiratory exchange rate, including heat expenditure, prior to termination. Body weight was recorded weekly, and the percentage change from baseline was calculated. In male mice, those maintained on a control diet (NPD-CM and PD-CM) showed a steady weight gain of approximately 35% from baseline, whereas HFD-fed males (NPD-HM and PD-HM) showed nearly ∼60% increase, plateauing at 12 weeks of feeding. In contrast, control diet-fed female mice (NPD-CF and PD-CF) demonstrated around 40% weight gain, while females on HFD showed a markedly higher increase of 75-90% with PD-HF mice reaching the highest levels (∼ 90%) and stabilising around 20 weeks. (Fig. 2a). Overall, all eight groups demonstrated gradual weight gain, with substantially higher increases in HFD-fed groups compared to their respective controls, highlighting the metabolic impact of prolonged high-fat feeding.

**Fig. 2.**
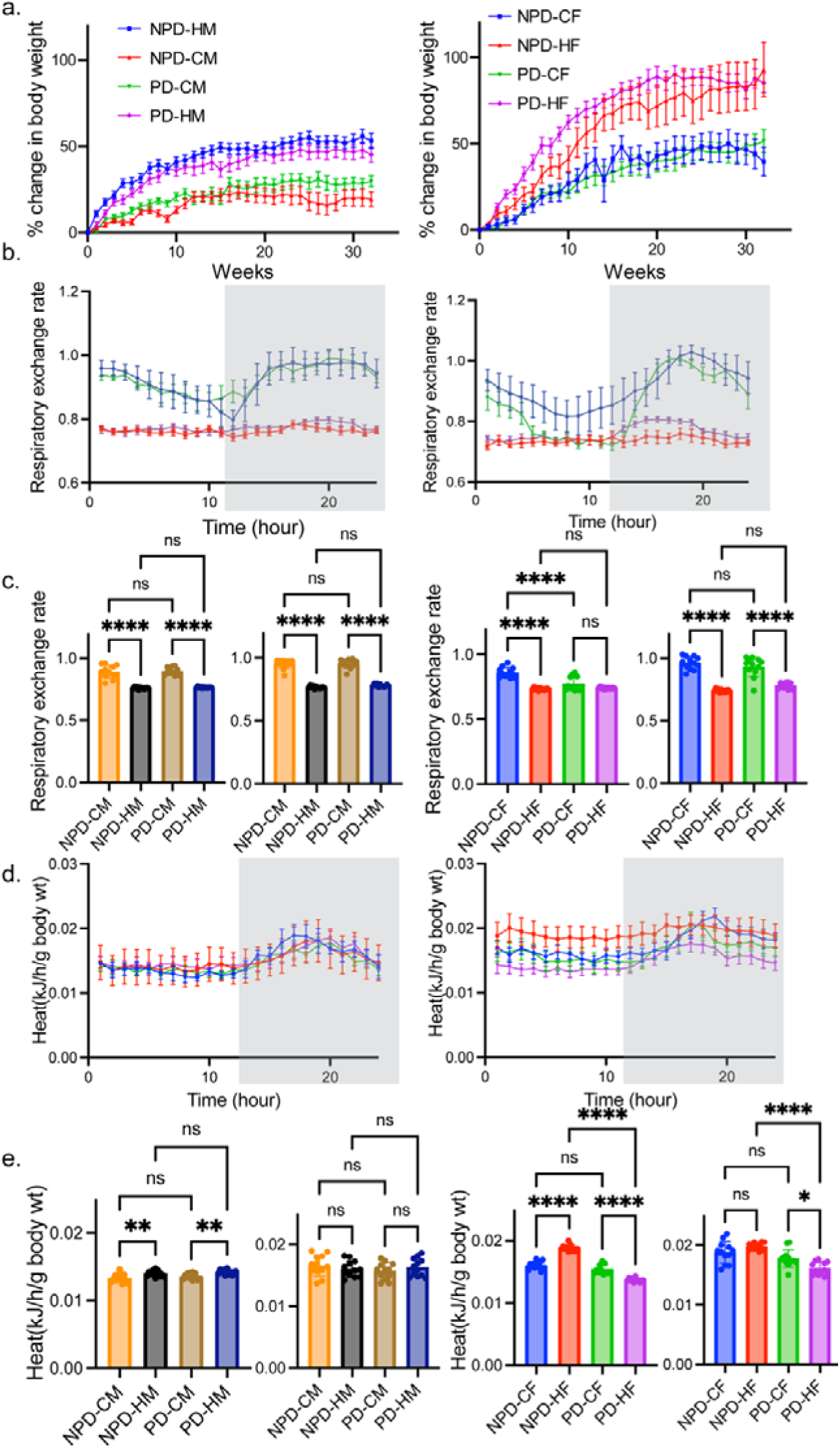
Effect of consumption of a high-fat diet on body weight and metabolism. a. body weight was measured every week from the start of feeding till euthanisation, and changes in body weight with respect to its starting weight were calculated using the formula (weight at that time /weight at starting *100-100). The graph on the left shows changes in the body weight of males (n = 13 for NPD-HM, n = 14 for NPD-CM, n = 13 for PD-CM, n=17 PD-HM) while the graph in the right represents changes in female body weight(n = 13 for NPD-CF n = 12 for NPD-HF, n = 16 for PD-CF, n = 15 for PD-HF). b. Respiratory exchange rate (RER) indicates utilization of fat as primary fuel in high-fat diet groups(H) while carbohydrate in the control diet group (C), the light-dark panel in the graph indicates the day-nighttime of the study. Graph in left shows RER pattern of male (n = 7 for NPD-HM n = 4 for NPD-CM, n = 5 for PD-CM, n=7 PD-HM) while graph in right represent RER pattern of female (n = 6 for NPD-CF n = 6 for NPD-HF, n = 6 for PD-CF, n=8 PD-HF). c. Cumulative representation of RER in Day(left) and night time(right) in both male and female. d. Energy expenditure is calculated in terms of heat produced by per unit weight in per unit time. Graph in left shows energy expenditure of male (n = 8 for NPD-HM n = 4 for NPD-CM, n = 5 for PD-CM, n=7 PD-HM) while graph in right indicates energy expenditure of female (n = 6 for NPD-CF n = 5 for NPD-HF, n = 8 for PD-CF, n=8 PD-HF). e. Cumulative representation of heat produced by per unit of body weight per unit time in Day(left) and night time(right) in both male and female Data are represented as mean ± SEM. * p < 0.05.

Next, we measured the respiratory exchange ratio (RER) to determine the predominant macronutrient being utilised by mice as an energy source during the day and night (active) phases. In all four control diet-fed groups, RER gradually declined during the daytime to an average value of ∼0.9, while remaining close to ∼1.0 during the night, indicating reliance on carbohydrate metabolism as the primary energy source. In contrast, all HFD-fed groups maintained a consistently lower RER ∼ 0.7 across both light and dark cycles, which suggests a metabolic shift towards fat utilisation as the primary fuel substrate. (Fig. 2b and c).

We further assessed heat production, which reflects energy expenditure relative to fat-free body mass. The heat produced per unit body weight in male mice was approximately 0.016 kJ/h across all four groups during nighttime. During the daytime, however, heat production was elevated by ∼5% in the NPD-HM group and 10 % in the PD-HM group compared with their respective control diet counterparts (NPD-CM and PD-CM). In females, daytime energy expenditure increased by 18% in NPD-HF mice relative to NPD-CF, with no significant difference observed during the night. Conversely, the PD-HF group exhibited a 12% and 9% reduction in energy expenditure during the day and night, respectively, compared to the PD-CF group. This distinct trend in female PD groups, characterised by reduced energy expenditure in PD-HF mice, likely contributed to the pronounced body-weight gain observed in these mice (Fig. 2d and e).

### High-fat diet-fed female transgenic mice exhibit earlier Parkinsonism than males

Hemizygous A53T transgenic mice typically develop Parkinsonian symptoms around 18 months of age^22^. HFD feeding in mice is known to accelerate disease progression^23^; however, its sex-specific impact has not been studied. To address this gap, we evaluated PD-associated phenotypic changes, including gut motility, anxiety, depression, and motor coordination in both male and female mice. Behavioural assays, including rotarod and grip strength tests, were initiated at 5.5 months of age and repeated fortnightly until mice showed any noticeable phenotypes. We assessed gut motility using the gold-standard carmine dye test, along with fecal output and fecal water content measurements. In both sexes, the time taken for the appearance of red feces was ∼2-fold longer in PD mice than in the NPD controls, indicating significant gut motility impairment. Notably, PD-HF exhibited ∼ 1.3-fold delay compared to PD-CF counterparts, and males showed no significant change (Fig. 3a). Furthermore, PD-HF displayed ∼2-fold reduction in fecal output and a 1.5-fold decrease in fecal water content compared to PD-CF mice, further supporting impaired gut motility in this group (Fig. 3b-d). We next evaluated anxiety and depression, two key non-motor symptoms of PD, in both sexes. Anxiety-like behaviour was measured through activity counts recorded in the Oxi-CLAMS activity counts compared with PD-CF mice, indicative of hyperactivity and increased anxiety-like behaviour (Fig. 3e–h). We assessed depression-like behaviours using the tail suspension and forced swim tests. Male groups did not show any significant differences, while PD-HF females spent ∼1.4-fold and ∼1.7-fold less active time in the tail suspension and forced swim tests, respectively, consistent with a depressive phenotype. To note, NPD-HF also exhibited reduced activity compared to NPD-CF, suggesting that HFD alone induces depressive-like behaviours in females, although the effect was more pronounced in transgenic mice (Fig. 3i and j).

**Fig. 3.**
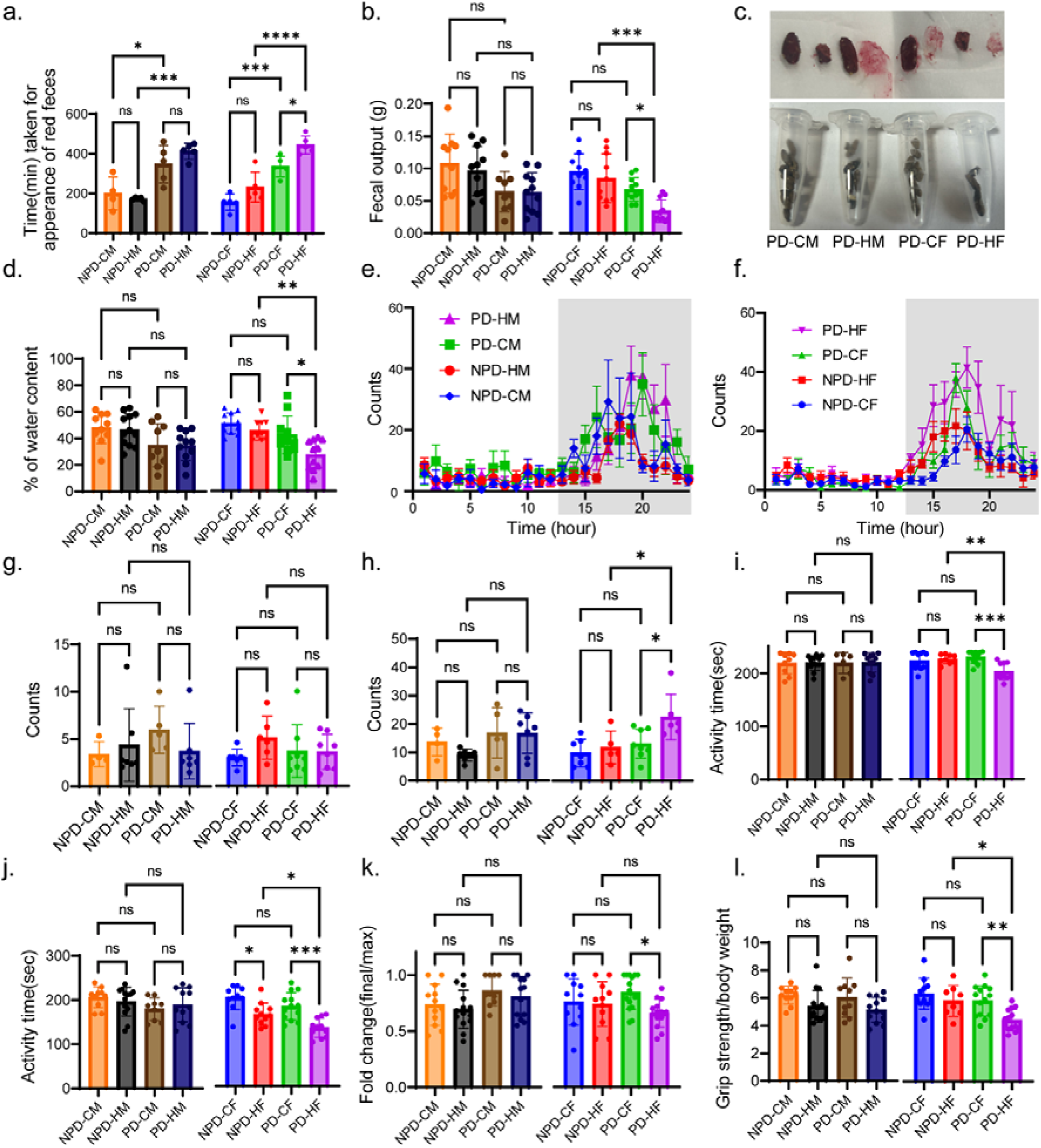
Phenotypic and behavioural assessment indicating early disease progression in females under the influence of HFD. **a.** Time taken for the appearance of first red colouration in faeces after the gavage of carmine dye(n = 5 for NPD-HM n = 4 for NPD-CM, n = 5 for PD-CM, n=5 PD-HM, n = 5 for NPD-CF n = 5 for NPD-HF, n = 5 for PD-CF, n=5 PD-HF) b. Weight of fecal pellets produced in one hour after the feeding of 2 hours (n = 11 for NPD-HM n = 10 for NPD-CM, n = 9 for PD-CM, n=11 PD-HM, n = 10 for NPD-CF n = 10 for NPD-HF, n = 11 for PD-CF, n=11 PD-HF) c. representative images of the amount of fecal matter collected and the red coloured faeces obtained in the carmine test. d. Percentage of water content in present in fecal matter (calculated by subtracting dry weight from the wet weight of feces) ((n = 12 for NPD-HM, n = 10 for NPD-CM, n = 9 for PD-CM, n=11 PD-HM, n = 11 for NPD-CF, n = 10 for NPD-HF, n = 11 for PD-CF, n=11 PD-HF)) e,f. Average counts moved per hour by a mouse in x-ampullary direction during 24 24-hour cycle, the light-dark panel in the graph indicates day and night time.( (n =7 for NPD-HM n = 4 for NPD-CM, n = 5 for PD-CM, n=7 PD-HM, n = 7 for NPD-CF n = 5 for NPD-HF, n = 7 for PD-CF, n=6 PD-HF) g,h. Average counts moved in daytime and nighttime, respectively. i. Time for which mice were in active state out of 240 sec of total duration of tail suspension test (n =12 for NPD-HM n = 9 for NPD-CM, n = 6 for PD-CM, n = 11 for PD-HM, n = 12 for NPD-CF n = 9 for NPD-HF, n = 11 for PD-CF, n=10 for PD-HF) j. Time for which mice were in active state out of total effective time of 240s during of Force swim test(n =13 for NPD-HM n = 10 for NPD-CM, n = 9 for PD-CM, n = 11 for PD-HM, n = 10 for NPD-CF n = 11 for NPD-HF, n = 13 for PD-CF, n=11 PD-HF) k. Ability of mice maintained for staying on moving rod as compared to their maximum ability, calculated in the form of fold change (final time taken for being on rod/ max duration for which mice could stay)( n =14 for NPD-HM n = 12 for NPD-CM, n = 10 for PD-CM, n=13 PD-HM, n = 11 for NPD-CF n = 11 for NPD-HF, n = 15 for PD-CF, n=14 PD-HF) l. Strength of mice to grip the grill per unit weight of body(n =14 for NPD-HM n = 11 for NPD-CM, n = 10 for PD-CM, n=13 PD-HM, n = 11 for NPD-CF n = 8 for NPD-HF, n = 14 for PD-CF, n=14 PD-HF) Data are represented as mean ± SEM. * p < 0.05.

We monitored motor coordination beginning 2 months after initiating dietary intervention and conducted fortnightly assessments using the rotarod and grip strength tests. Female PD-HF mice showed a gradual decline in rotarod performance from week 15, which became significant by week 21. In contrast, PD-HM males showed no significant change up to week 32 (Fig.S 1a). Grip strength in PD-HF also declined gradually, becoming evident from week 32 onward (Fig.S1b). At study termination, we calculated the fold change in rotarod retention time (relative to maximum performance) and grip strength normalised to per body. PD-HF demonstrated ∼1.25-fold and ∼1.6-fold reductions in rotarod performance and grip strength, respectively, confirming motor coordination deficits in this group (Fig. 3k and l).

### Molecular changes in ENS and CNS support early disease progression in females

Phenotypic assessments indicated early PD progression in the PD-HF group. According to Braak’s hypothesis, PD pathology is initiated in the gut and subsequently spreads to the brain, with impaired gut motility serving as an early hallmark of the disease^24,25^ Since the enteric nervous system (ENS), often referred to as the “second brain,” is known to be affected in PD even before central nervous system (CNS) involvement, we analysed both ENS and CNS to evaluate molecular correlates of disease progression.

We assessed phosphorylated (p129S) α-synuclein (p-αSyn), a gold-standard marker of PD pathology, in both ENS and CNS, and measured tyrosine hydroxylase (TH) levels as an indicator of dopaminergic neuron integrity in CNS only. For ENS pathology, we stained the whole mounts of the muscularis layer containing the myenteric plexus with p-αSyn and UCHL1. We quantified the intensity of p-αSyn normalised to UCHL1 and found that PD groups displayed ∼3.5-fold higher intensity compared to NPD groups. Within PD groups, PD-HF mice exhibited a ∼1.5-fold increase compared to PD-CF, whereas PD-HM did not differ significantly from PD-CM counterparts (Fig. 4a and b).

**Fig. 4.**
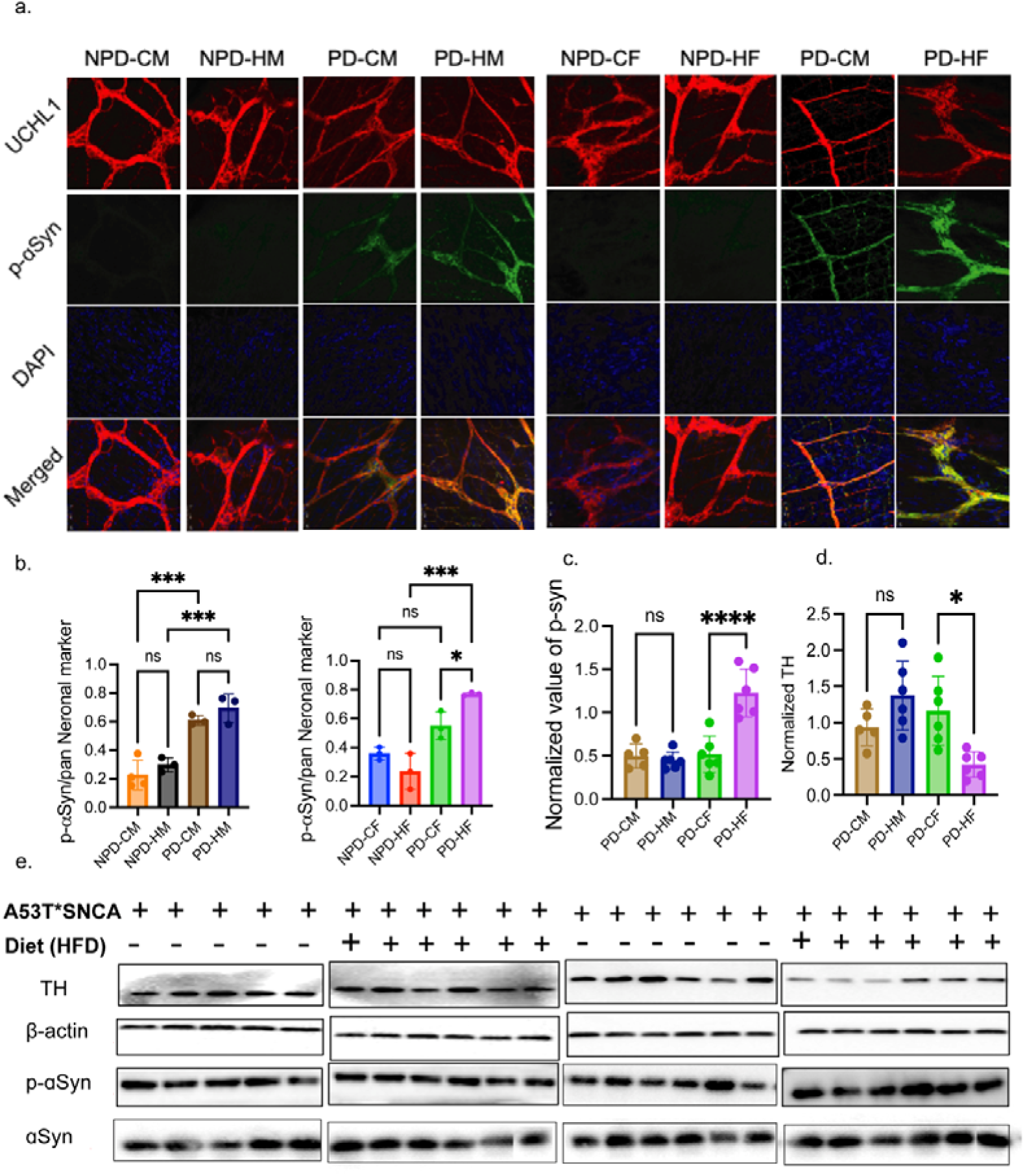
Change in the level of PD markers in the Enteric and Central Nervous System. a. Representative panel ( left: male and right female) indicating changes in the level of p-αSyn in enteric neurons, UCHL1(red) act as neuronal marker while DAPI( blue) is used as nuclear stain (N=3 with three regions in each mice). b. Quantification of the microscopic data c. d. Quantification of western blot data of p-αSyn and TH respectively. e. Representative western blot of brain lysate indicating changes in the level of Tyrosine hydroxylase(TH), a marker of dopaminergic neurons and p-αSyn. Β-actin is used as a reference protein.

In CNS tissue, we evaluated αSyn and p-αSyn levels by western blot analysis. The NPD groups displayed negligible αSyn expression, with no quantifiable p-αSyn bands. In contrast, PD groups showed clear and quantifiable αSyn and p-αSyn bands (Fig. 4e). When normalised p-αSyn levels were compared, only PD-HF mice demonstrated a ∼2-fold increase over PD-CF, with no significant difference between PD-HM and PD-CM (Fig. 4c and e). In addition, TH levels were significantly reduced in PD-HF mice, further confirming early and accelerated disease progression in this group (Fig. 4d and e).

### Proteomics data reveal sex-dimorphic changes associated with PD

To investigate the molecular basis of sex-dimorphic patterns in PD progression under HFD conditions, we employed a mass spectrometry (MS)-based proteomics approach. We analysed whole-brain lysates using a data-independent acquisition (DIA) workflow for quantitative protein profiling. Approximately 3,200 proteins were identified per sample (Fig. 5a). We validated the data quality by searching all samples in a single combined file. Principal component analysis (PCA) clustered the dataset in PC1 (18.63% variance) and PC2 (28.93% variance), revealing a clear segregation between PD and NPD groups, with minor overlap of the PD-CF group into the NPD clusters. However, we did not observe distinct clustering based on sex (male vs. female) or diet (control vs. high fat) within either NPD or PD groups (Fig. 5b). The average median coefficient of variation (CV) across precursor ions was ∼28%, supporting that group separations represent true biological signatures of altered proteomes (Fig. 5c). After confirming data quality, we conducted a series of multiple comparative analyses. First, we evaluated the effect of a single copy of the A53T SNCA mutation in a sex-specific manner by comparing PD-CM vs. NPD-CM and PD-CF vs. NPD-CF. We defined differentially expressed proteins (DEPs) using a threshold fold change ≥1.4 for upregulation and ≤0.714 for downregulation with FDR ≤0.05. In males (PD-CM vs. NPD-CM), 154 proteins were downregulated and 40 upregulated (Fig. 5d). In females (PD-CF vs. NPD-CF), 87 proteins were downregulated and 92 upregulated (Fig. 5e). We next performed pathway enrichment analysis of these DEPs to uncover biological processes affected by the mutation. Both sexes showed negative enrichment of mitochondrial pathways, including Complex I biogenesis, respiratory electron transport, ATP synthesis by chemiosmotic coupling, uncoupling protein-mediated heat production, and the TCA cycle, consistent with mitochondrial dysfunction as a central mechanism in PD. (Fig.S2a).

**Fig. 5.**
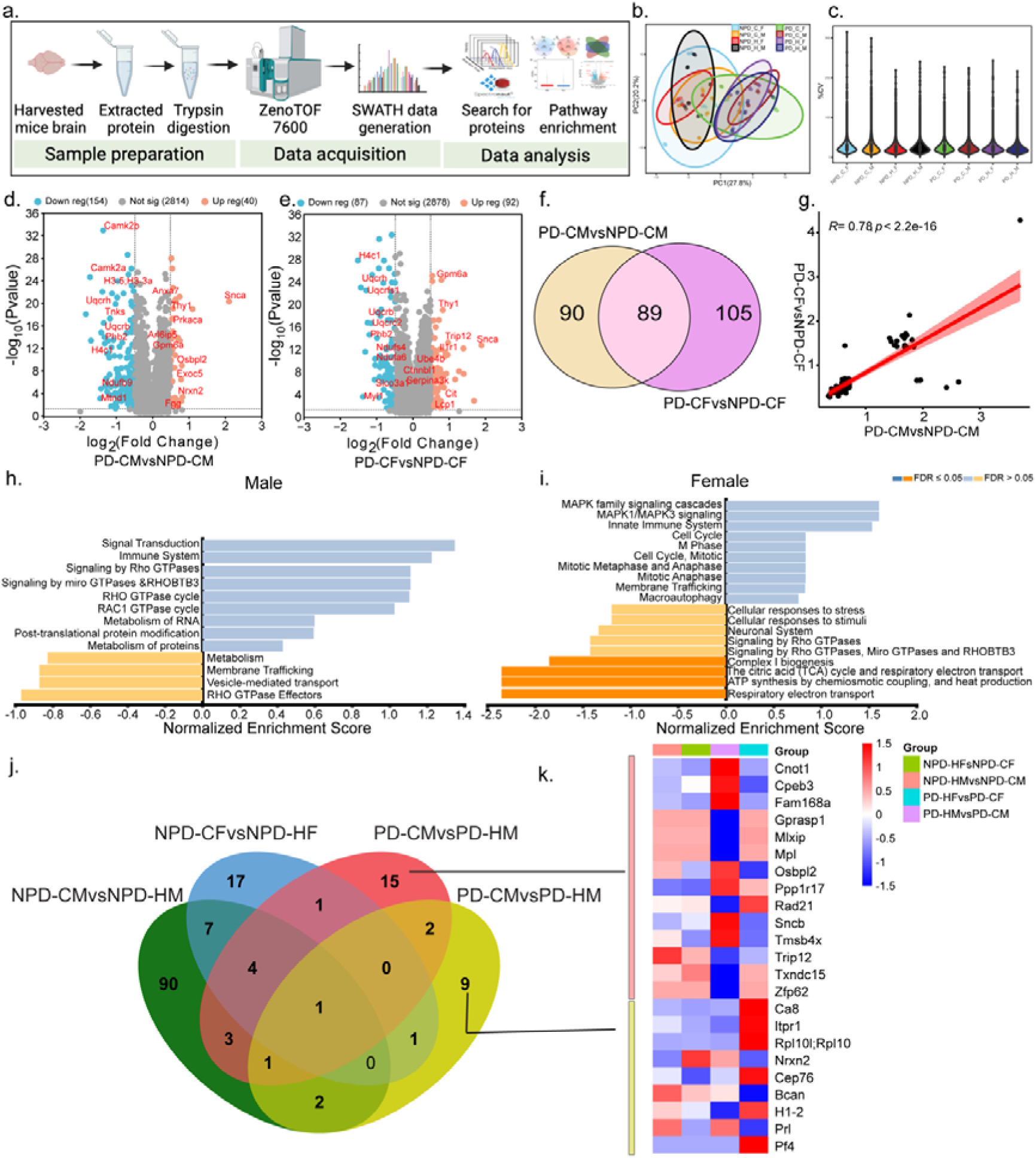
Sex specific effect of incorporation of A53T mutant form of. α**Syn** a. schematic representation of workflow used for sample preparation , data acquisition and data analysis b. Principal component analysis of samples clustering based on their comparison groups separated by PC1(28.93%) on X-axis and PC2(18.63%) on Y-axis. Effect of genotype is clearly evident on clustering pattern of PD and NPD groups. While no clustering is present based on biological sex and diet administered is observed. C. Percentage CV distribution of identified precursors among the samples from different comparison group is approximately similar(26%). d. Volcano plot for the comparison between PD and NPD male mice groups, feed on control diet, indicating total identified protein along with differentially regulated proteins(DEPs) , x-axis in represent log2 fold change and Y-axis represents. e. Volcano plot for the comparison between PD and NPD female mice groups, feed on control diet, indicating total identified protein along with differentially regulated proteins(DEPs) f. Venn diagram representing common and specific DEPS identified in male and females when compared between PD and NPD groups g. Correlation plot indicating that common DEPs are following same trend of expression, X-axis represent fold change of protein in male and Y-axis indicated fold change in female h. Pathways enriched using the fold change of male specific DEPs indicating 9 positively enriched and 4 negatively enriched pathways. X -axis represents normalized enrichment score of the pathways. All belongs to FDR> 0.05. i. Pathways enriched using the fold change of female-specific DEPs, indicating 10 positively enriched and 9 negatively enriched pathways, out of which 4 belong to FDR ≤ 0.05. j. Venn diagram of DEBs of DEPs when compared high fat administered group with the control diet administered group by keeping genetic background and biological sex constant; NPD-HMvsNPD-CM, NPD-HFvsNPD-CF, PD-HMvsPD-CM, PD-HFvsPD-CF. j Heat map indicating fold change of PD-HMvsPD-CM specific and PD-HFvsPD-CF specific DEPs among all 4 comparisions.

To identify sex-specific alterations, we compared the DEPs from PD-CM vs NPD-CM with those from PD-CF vs NPD-CF. We found 89 DEPs common to both sexes, while 90 were unique to males and 105 were unique to females (Fig. 5f). The common DEPs showed positive correlation (r = 0.78), indicating shared proteomic alterations across sexes (Fig. 5g). Gene set enrichment pathway analysis of unique DEPs revealed male-specific enrichment of pathways, including Rho and Miro GTPase signaling, RHOBTB3 signaling, post-translational protein modification, metabolism, and membrane trafficking. In contrast, female-specific DEPs pathways enriched MAPK family signalling, cell cycle, regulation, macroautophagy, and TCA cycle (Fig. 5h and i).

Next, we evaluated the effect of diet as a variable under genetic and sex specific conditions. We performed four pairwise comparisons, including NPD-HM vs. NPD-CM, NPD-HF vs. NPD-CF, PD-HM vs. PD-CM, and PD-HF vs. PD-CF, which yielded 108 DEPs (11 downregulated, 97 upregulated), 31 DEPs (13 down, 18 up), 27 DEPs (19 down, 8 up), and 31 DEPs (4 down, 12 up), respectively, using the same criteria of fold change ≥1.4 and ≤0.714 with FDR ≤0.05 (Fig.S 2b-e). When we compared all four datasets, we found 90 DEPs unique to NPD-HM vs. NPD-CM, 17 unique to NPD-HF vs. NPD-CF, 15 unique in PD-HM vs. PD-CM, and 9 unique in PD-HF vs. PD-CF (Fig. 5j).

Notably, the unique DEPs identified in PD-HM vs. PD-CM included upregulation of CCR4-NOT transcription complex subunit 1 (Cnot-1) and synuclein beta (Sncb), both of which have been previously documented to exert neuroprotective effects. In contrast, unique DEPs in PD-HF vs. PD-CF included upregulation of carbonic anhydrase 8 (Car8) and downregulation of neurexin II (Nrxn2), suggesting synaptic dysfunction as a female-specific molecular signature.

### Alteration in synaptic proteins as a potential driver of early disease progression in females

After analysing each variable individually, we next evaluated the cumulative effect of diet and the A53T mutation. We compared PD-HM vs. NPD-HM and PD-HF vs. NPD-HF. The PD-HM vs. NPD-HM and identified 184 downregulated and 25 upregulated proteins, while PD-HF vs. NPD-HF displayed 154 downregulated and 48 upregulated proteins (Fig. 6a,b).

**Fig. 6.**
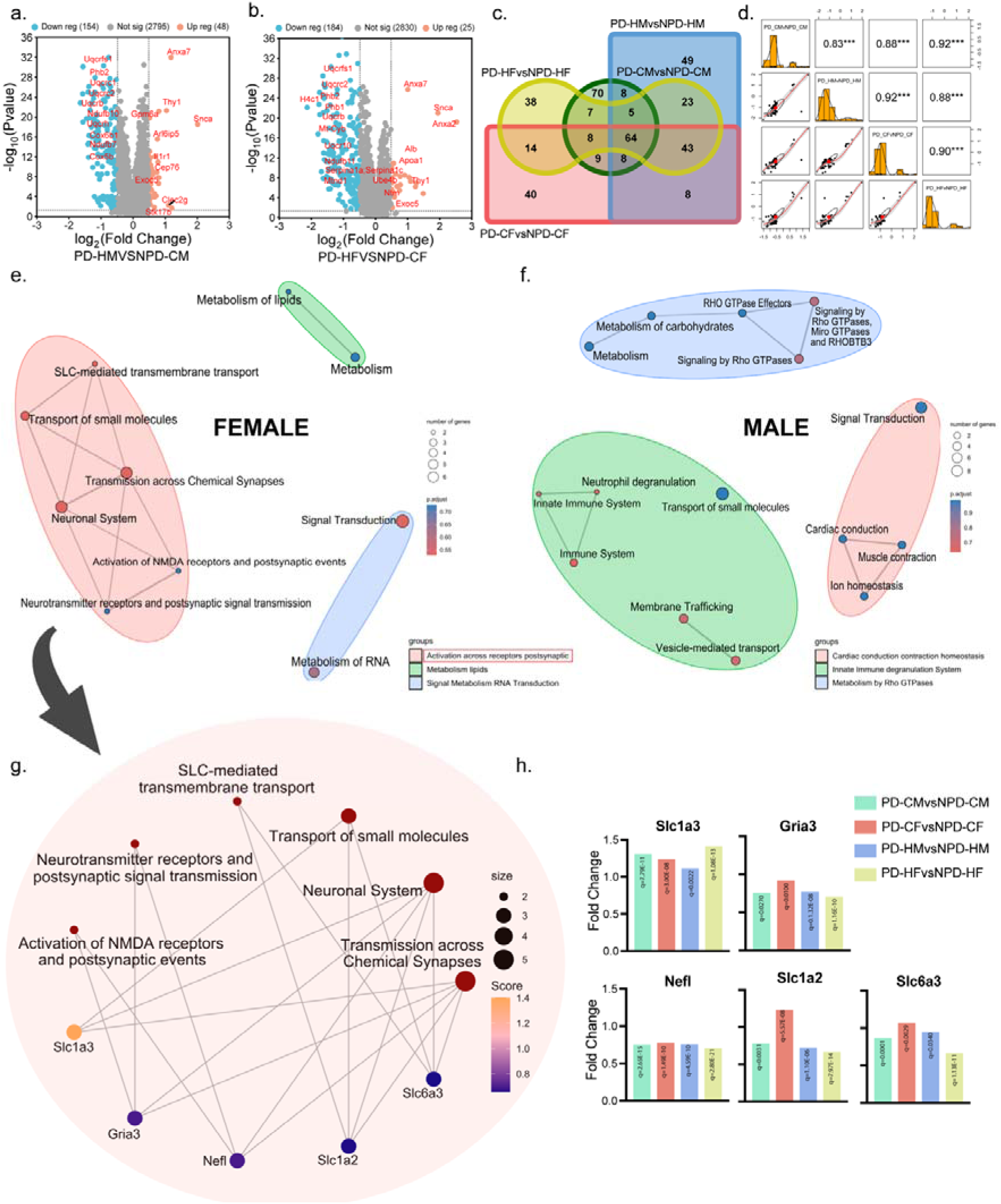
Sex-specific pathways and their associated proteins indicate differential modes of disease progression. a. Volcano plot indicating identified proteins and DEPS on comparison between PD and NPD groups when fed on a high-fat diet to study the cumulative effect of HFD and gene in the male group. b. Volcano plot indicating identified proteins and DEPs on comparison between female PD and NPD group when fed on a high fat diet. c. venn diagram showing common and specific DEPs (PD vs NPD) among four comparisons d. representation of correlation present among the fold change of common DEPs from the four comparisons e. cluster of pathways enriched using Male_HFD specific DEPs f. cluster of pathways enriched using female HFD specific DEPs g.DEPs from the female specific pathway, along with their normalized intensities acquired from MS.

Pathway enrichment of these DEPs revealed a negative enrichment of mitochondrial-related pathways, including Complex I biogenesis, metabolism, ATP synthesis by chemiosmotic coupling, the TCA cycle, and respiratory electron transport in both sexes. Conversely, we observed a positive enrichment of immune-related pathways such as cytokine signaling and RNA metabolism. In male mice, enriched pathways included vesicle-mediated transport, neutrophil degranulation, interferon signalling, post-translational protein modification, and protein metabolism. In contrast, females showed enrichment of pathways associated with MAPK signalling, neurotransmitter receptor and postsynaptic signal transmission, and ion and amino acid transport (Fig.S2f).

To better understand these patterns, we compared DEPs from PD-HM vs. NPD-HM, PD-HF vs. NPD-HF, PD-CM vs. NPD-CM, and PD-CF vs. NPD-CF using a Venn diagram and correlation plots (Fig. 6c and d). We observed high correlations (r = 0.83-0.92) among the common DEPs, suggesting consistent regulatory trends across different groups. Gene set enrichment analysis of unique DEPs revealed male-specific clustering of pathways related to cardiac immune conduction, membrane signaling, small molecule metabolism, and Rho GTPase signaling. In contrast, female-specific clusters included postsynaptic receptor activation, RNA metabolism, and lipid signaling transduction (Fig. 6e and f). Among these, the postsynaptic receptor activation cluster contained the greatest number of pathways, suggesting a central role in the observed phenotypic and molecular changes. Key pathways in this cluster included neurotransmitter receptor and postsynaptic signal transmission, chemical synapse activity, NMDA receptor activation, postsynaptic events, and SLC-mediated transmembrane transport. The corresponding altered proteins included Gria3, Slc1a3, Slc1a2, Slc6a3, and Nefl (Fig. 6g). When we calculated the fold change of these proteins in PD groups with respect to their NPD counterparts, we observed a significant decrease (FC ≤0.714) in the levels of Gria3, Slc1a2, Nefl, and Slc6a3, with an increased level of Slc1a3 (FC=1.403) in the PD-HF group. (Fig. 6h). We further validated the expression profile of Gria3 and Slc1a2, along with scaffolding protein Homer-1, using western blot, which confirmed a significant reduction in synaptic protein levels. (Fig. 7a-d)

**Fig. 7.**
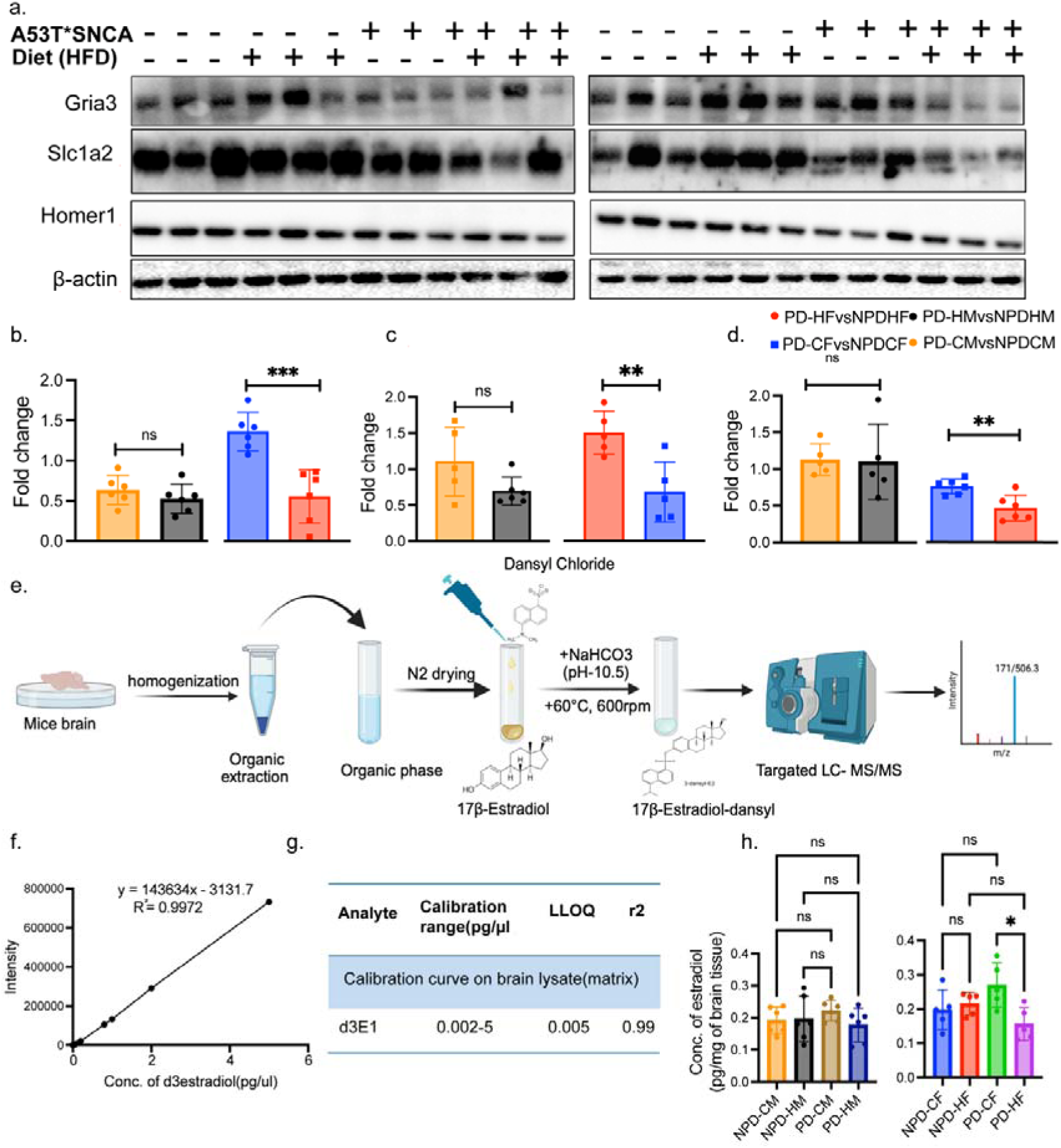
Reduced synapse-associated protein and estradiol level in females under the influence of high-fat diet. **a.** Representative western blot of brain lysate indicates decreased synapse-associated DEPs, Gria3 and Slc1a2, along with HOME1, a synaptic scaffold protein, in the PD-HF group. b.c.d. Quantifications of fold change in PD-high fat groups compared to NPD-high fat from western blot data. e. Scheme used for the extraction and derivatisation of estradiol from the brain tissue, along with the representative MS spectra achieved from targeted MS/MS. F. Standard curve, for estimation of estradiol level in brain tissue, prepared using linear concentration of d3E1 on matrix(brain tissue) g. Table indicating calibration range, LLOQ and r2 d3E1 used for standard curve preparation. h Concentration of estradiol in Pg per unit brain tissue, indicating a significant decrease in the level of estradiol in the PD-HF group.

**Fig. 8.**
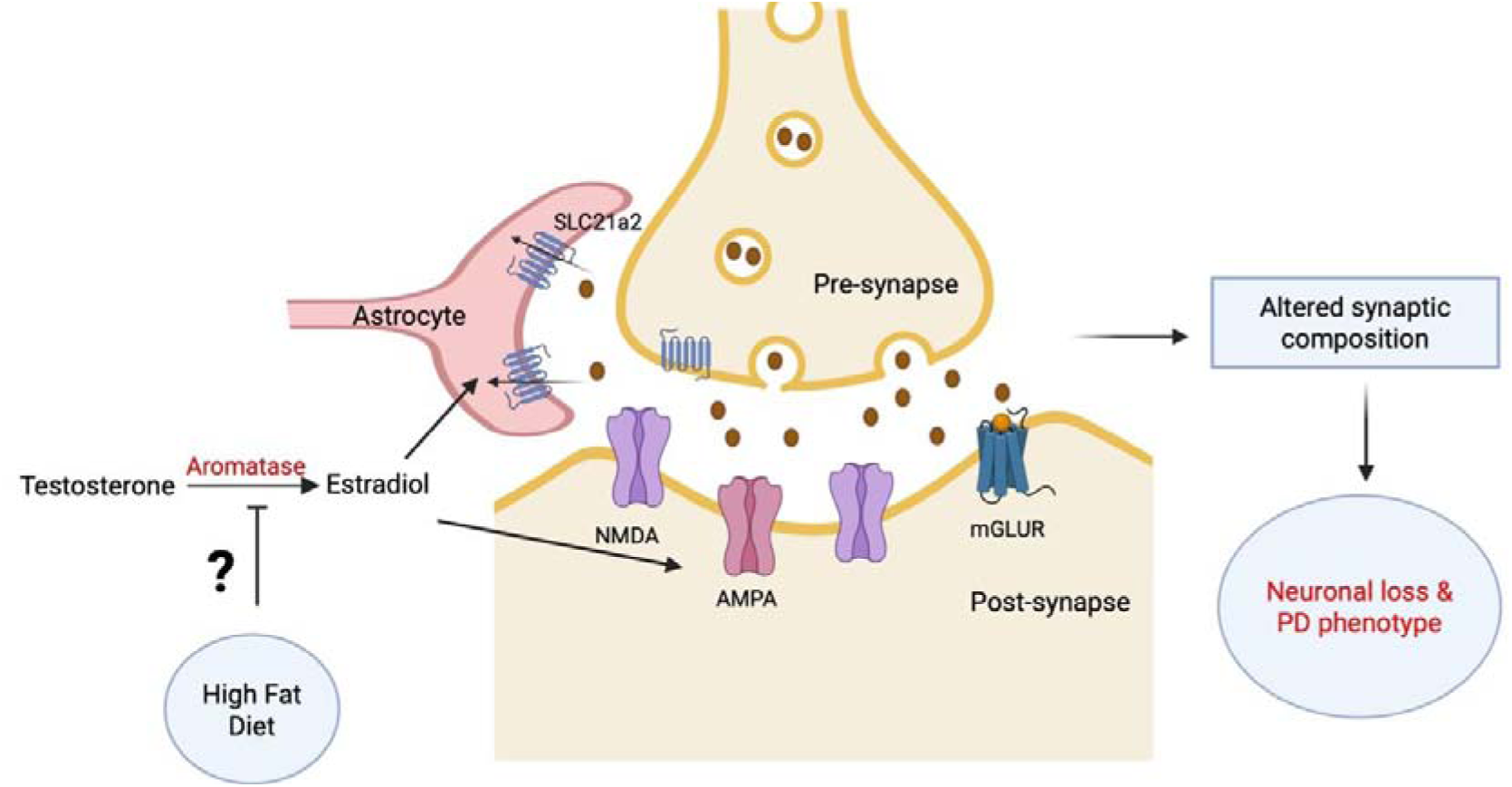
Suggestive mechanism of early PD progression in females under the influence of HFD. Illustration indicating that HFD is affecting the estradiol synthesis mechanism, leading to decreased levels of estradiol in the brain, which is necessary to maintain the composition of various ion channels at synapses. This alteration in synaptic protein levels affects the synaptic plasticity and associated transmission, which in turn leads to neuronal loss with visible PD symptoms.

### Lack of neuroprotection from estradiol as a potential cause of synaptic alterations

Estradiol, a female sex hormone, is well recognised for its neuroprotective role and its ability to reduce the risk of PD in reproductively active females. One of its key functions is the maintenance and enhancement of synaptic strength in brain regions. Since we observed alteration in the synaptic proteins in the PD-HF group of our study, we evaluated estradiol levels to better understand its potential contribution to early disease progression under HFD conditions in females. We quantified estradiol after derivatisation with dansyl chloride and quantified the intensity of the resulting adduct using targeted MS (Fig. 7e). We generated a standard calibration curve using the deuterated (d[) form of estradiol across a concentration range of 0.002–5.00 pg/µl, using brain lysate as the matrix (Fig. 7f and g). The measured estradiol concentrations in brain tissue ranged from 0.11 to 0.36 pg/mg. While no significant sex differences were observed between male and female groups overall, with a marked reduction in the PD-HF group compared to PD-CF, reflecting weaker estradiol-associated neuroprotection in females under HFD conditions (Fig. 7h). This loss of estradiol may underlie the observed synaptic dysregulation and accelerated disease progression in females.

## Discussion

This study demonstrates that PD progression occurs in a sex-dependent manner in the A53T α-synuclein transgenic mouse model, with female mice showing earlier onset of behavioural and molecular symptoms than males. These symptoms included anxiety, depression, impaired gut motility, and reduced motor coordination, all of which were exacerbated under HFD conditions. The observed increase in body weight and altered respiratory exchange ratio confirm the metabolic effects of a high-fat diet in both PD and NPD groups. A similar effect is reported in other PD models, particularly in males, where chronic fat intake leads to obesity and metabolic imbalance^12^. However, few studies have also described either resistance to weight gain or reversal of the HFD effect upon diet withdrawal^20,26^. The alterations in gut motility, anxiety, and depression like behaviours are the early non-motor features of PD, whereas motor impairment and loss of muscle strength emerge as late signs of the disease^27,28^. In our study, male mice displayed decreased gut motility and increased anxiety without noticeable changes in motor coordination, indicating an early stage of disease and a limited impact of HFD. On the other hand, HFD-fed female PD mice exhibited a significant decrease in gut motility, increased anxiety and depression, along with loss of motor coordination and grip strength consistent with the late stage of the disease. Increased p-αSyn accumulation in both ENS and CNS, accompanied by reduced TH levels, further confirms accelerated neuropathology. These findings align with previous evidence showing that female mice display comparable vulnerability to HFD in models of Alzheimer’s disease and mixed dementia^29^. Moreover, recent studies have reported that women with diabetes, prediabetic conditions, high-fat food consumption, and obesity are at increased risk of developing PD^30,31^. The outcomes of our animal study corroborate these clinical observations, strengthening the link between metabolic dysfunction, dietary fat intake, and heightened susceptibility to PD in case of females.

The proteomics studies on the mouse brain revealed enrichment of pathways related to the respiratory electron transport chain, TCA cycle, complex I biogenesis, and metabolism in both PD-CM and PD-CF groups compared to their NPD counterparts, indicating mitochondrial dysfunction, a hallmark of early PD pathology in both sexes^32,33^. In addition, male-specific immune-related pathways like cytokine signalling and neutrophil degranulation, and female-specific pathways like pathways related to rRNA processing, nonsense-mediated decay, and ion channel transport highlight the sex specific effect of A53T*SNCA transgene. We next evaluated the combined effect of HFD and A53T incorporation by comparing PD-HM and PD-HF with their respective NPD counterparts. This analysis revealed several PD-related pathways common to those observed under control diet conditions, along with a subset of sex-specific pathway alterations unique to each group. In males, pathways related to the immune regulation and metabolic function were primarily affected, consistent with the previous report^34^. To the best of our knowledge, there are no prior studies that have addressed female-specific molecular responses to HFD in PD models.

Our data demonstrate that in females, signal transduction, metabolic, and particularly synapse-associated pathways are affected, while immune responses remained relatively unaltered.

The association between synaptic dysfunction and PD progression is well documented in the literature, and our findings suggest that synaptic vulnerability may be a potential cause of early disease progression in females under the influence of HFD^35,36^. Supporting this, we observed decreased levels of Nrxn2, a key protein regulating synaptic junction integrity and previously implicated in PD pathology, in the PD-HF as compared to the PD-CF^37,38^. Furthermore, we quantified the level of β-estradiol, a female-specific hormone known for its neuroprotective role in maintaining synaptic function^39^.β-Estradiol levels were significantly reduced in the PD-HF group, indicating loss of estradiol-mediated synaptic protection. This likely contributes to synaptic degeneration and early disease progression in PD-prone females under HFD^40,41^

In summary, this study demonstrates that female A53T α-synuclein transgenic mice develop an accelerated Parkinsonian phenotype when fed with high-fat diet. The early loss of synaptic function likely contributes to faster disease progression in females. The female sex hormone estradiol provides neuroprotection by maintaining synaptic integrity in normal dietary conditions. However, a high-fat diet may disrupt estrogen biosynthesis and signalling, leading to reduced estradiol levels and subsequent synaptic vulnerability. Future studies will investigate how HFD-induced metabolic stress and estradiol loss contribute to synaptic dysfunction in Parkinson’s disease.

### Methods Mice

Two breeding pairs (1:2) of hemizygous (containing a single copy of SNCA*A53T with Prnp promoter) B6;C3-Tg (Prnp-SNCA*A53T)83Vle/J (strain #004479) were purchased from Jackson Laboratory, USA. From the F1 generation, multiple breeding between hemizygous pairs (1:2) were set up to produce hemizygous and heterozygous mice at the animal breeding facility (RCB, Faridabad). Mice were housed in groups of up to 5 in individually ventilated cages with a 12 hours light-dark cycle, and food and water were provided *ad libitum.* From the age of 3.5 months, both male and female mice were incorporated for further studies. All animal experiments were approved by the Regional Centre for Biotechnology Institutional Animal Ethics Committee (IAEC Animal Use Protocol #2021-0162)

### Metabolic Assessments

For assessing metabolic changes, including changes in substrate utilization, energy expenditure (heat), VO2, and VCO2, an eight-chambered Oxymax-Comprehensive Lab Animal Monitoring System (Oxi-CLAMS) was used. Cages were cleaned and autoclaved prior to the initiation of the experiment. Mice were acclimatized for 12 hours, followed by the assessment of all the parameters for the next 24 hours. Food and water were provided *ad libitum*.

### Body weight and food intake

Body weight was measured weakly between 10:00 am to 1:00 pm. Food intake was also measured weakly by weighing the food pellets left in the mice after giving a known amount of food.

### Gut motility test

To assess the motility of the gut, the gold standard test, the carmine dye test, along with the amount of stool passed and water content in feces, was performed. All the tests were conducted before initiation of final behaviour studies to avoid activity-mediated changes. To avoid day-night bias, all the tests were started from 10:00 am.

#### Fecal output and water content in feces

Animals were maintained in a fasting state for sixteen hours, followed by providing food *ad libitum* for 2 hours. Then each mouse was transferred to a clean cage, without bedding and food, for one hour. Water was provided *ad libitum* throughout the experiment. Fecal pellets were collected in individual pre-weighted MCTs. The weight of feces with an MCT was measured freshly and allowed to dry for 18 hours at 65°C, and the dry weight was taken. For the calculation of the percentage of water in faeces Percentage (%) of Water = (wet weight-dry weight)/wet weight *100 was used.

#### Carmine test

Carmine dye (6% w/v) was prepared in 0.5% methylcellulose solution and autoclaved before exposure to mice. No fasting was provided to the mice before the head. Mice was gavaged with 130 μl of carmine solution between 9:30 am to 11:00 am and kept singly in cages without bedding, provided food and water *ad libitum.* Feces were checked for red colouration at every 30 min interval, and the time for the appearance of red colour in faeces was recorded.

### Behavioural Studies

Several behavioural tests were performed to assess various parameters like motor coordination, strength of gripping, depression, and anxiety. All the mice were gently treated by the same individual throughout the experiment. Ten min prior to the start of the experiment, cages were opened to awaken and make the mice active. Only one or two tests were conducted in a single day to avoid the exhaustion of the mice. The platform of all the instruments was properly cleaned before the use of each animal. All the tests are performed between 10:00 am to 6:00 pm in the light-on cycle. Mice were exposed in randomized order to avoid the biasness.

#### Rotarod test

The Rotarod test was conducted using a rotarod apparatus (IITC-Lifescience Rotarod Series 8), which was programmed to rotate with a gradual increase in speed from 5 rpm to 40 rpm in 50 seconds. Magnetic sensors on the plate get activated when mice fall from the rod and give the reading of the time taken to fall. Mice were pre-trained for 3 consecutive days with three trials on each day, and provided a 3 min rest in between trials. The Rotarod test was initiated after 2 months of feeding and continued once every fortnight to assess the changes in motor coordination throughout the experiment. For experimental purposes, a maximum of three consecutive trials with 4 - 5 min rest in between were taken. To assess the progress of the disease at the end of the experiment, the fold change occurring in “Retention time” was calculated using Fold change = Time to fall at the end of the experiment/the maximum time to fall from the initiation of the experiment.

#### Tail suspension test

Mice were adhered to a surface 100 cm above the base by their tails, 2 cm away from the tail tip, with the help of adhesive tape. The activity of mice was recorded for 4 min and analysed by an experimentally blinded person. A maximum of 4 mice were adhered at a time, maintaining a minimum distance of 20 cm. Activity time was calculated by Activity time = Total time - immobility time (when there is no movement in any limb)

#### Force swim test

Mice were placed in a 5 L glass cylinder (40 cm high, 25 cm diameter) filled with water up to the 4 L mark. The temperature of water was kept constant for all mice (28 ± 2 °C). For assessing the behaviour, videography was done for 6 min. The initial 2 min were considered as acclimatisation time, while the last 4 min were considered for analysis. Immobility state was defined as a condition with a lack of movement of the forelimb with minimal movement in the hind limb, which is necessary to keep the head above. The analyst was kept blinded about the group of mice to avoid bias, and mice were placed in randomised order. Activity time was calculated by Activity time = Total time - immobility time (when there is no movement in any limb)

#### Grip Strength Test

For measuring grip strength, BIOSEB-GRIPTEST v3.48 instrument was used. Mice were kept on the grid and allowed to hold it for 1 - 2 sec, followed by pulling with the base of the tail with constant force. The strength of grip in grams was calculated by the instrument. To avoid the effect of body weight on grip strength, it was normalized with body weight.

### Proteomics sample preparation and analysis

Whole brain lysate was used for a proteomics study. The brain was minced thoroughly, and approximately 30 mg of tissue was dissolved in RIPA buffer. The tissue was homogenized using a hand homogenizer. Further cells were lysed using a probe sonicator, and the supernatant was used for protein quantification. 50 µg protein was taken from each sample and reduced and alkylated using 10 mM DTT for 30 min at 56 °C and 20 mM IAA for 1 h at RT. These reduced proteins were digested using 1:20 trypsin (Pierce, Thermofisher) at 37 °C for 18 h. The digested peptides were desalted using a C18 cartridge (Oasis HLB, Waters), and the eluted peptides were vacuum dried and stored at -80 ^0^C. The dried peptides were resuspended in 2% Acetonitrile in water in the presence of 0.1% formic acid prior to LC-MS injection.

Data acquisition was performed with a ZenoTOF 7600 (Sciex, MA, USA) mass spectrometer, coupled with a Waters ACQUITY UPLC M-Class System, and data were acquired in SWATH mode. An equivalent quantity of digested peptide (1µg) was loaded on Luna 5 µm C18 Micro trap column (20 mm × 0.3 mm, 100 Å, Phenomenex, Torrance, CA, USA) which was further resolved on nanoEase M/Z HSS T3 analytical column (150 mm × 300 µm, 1.8µm, 100 Å, Waters). Peptides were eluted through a linear gradient of acetonitrile (with 0.1% (v/v) formic acid) for 22 min, for a total run duration of 32 min, with a flow rate of 5 μL/min. An accumulation period of 20 ms with a cycle time of 1.74 sec was used on SWATH acquisition mode. A variable window was created to cover a mass range of 400-1250 Da, providing 65 variable-size overlapping windows.

The data were processed using Spectronaut Pulsar 18 (Biognosys), and a protein identification. A de novo library was generated in Spectronaut Pulsar 18 (Biognosys) using wiff files of all the samples, keeping the UniProtKB mouse protein database. This generated library was further used for protein identification in each sample. Global and dynamic parameters are employed for the normalization strategy and XIC RT extraction, respectively. The enzyme specificity is set for trypsin with a maximum missed cleavage of 2. For further analysis, the FDR value was kept at 0.05, and PGquant values were used as intensities. WebGStalt and ShinyGo were used in the downstream analysis of pathways, and the data representation was done using SRplot and R software (R 4.3.1).

### Western blotting

The western blotting experiments were performed with the standard protocol. Briefly, 12 μg protein was loaded for studying highly abundant proteins, while for less abundant proteins, 40 μg protein was loaded on a 1.5 mm gel. After electroporation, the gel was wet transferred for 75 min, followed by blocking using skimmed milk (5%). The blot was incubated in the primary antibody(Tyrosin hydroxylase(abcam,1:1000); p-αSyn(abclonal,1:1000); αSyn(CST,1:3000);Homer-1(Invitrogen,1:4000); Gria3(Invitrogen,1:1500); GLT-1(Invitrogen,1:3000); β-actin(abclonal,1:5000)) overnight at 4°C. Based on the requirement, the secondary antibody(anti-rabbit(Invitrogen,1:8000); anti-mouse(Invitrogen,1:8000)) was used for 2 hours at RT. Blots were visualised in ChemiDoc^TM^ XRS+ (Bio-Rad), and band intensity was measured using Image Lab software.

### Isolation of the myenteric plexus

The gut of the mice was isolated after euthanisation and flushed with PBS to remove the fecal matter. A sterile plastic stick was inserted into the lumen to enwrap it, and the outermost layer was peeled using a wet cotton swab placed on another stick. This layer was stretched on a glass slide and fixed with PFA (4% in PBS) for 20 min. After washing, it was stored at - 20°C until further use.

### Staining and microscopy

To study the changes in ENS, the whole mount of the muscular layer embedded with the myenteric plexus was used. A section of the whole mount was taken off the slide and stained using the free-floating technique. Briefly, the section was antigen-retrieved using citrate buffer (pH 8.0) at 95°C for 30 min. After washing, it was blocked with blocking solution (5% BSA, 5% FBS, 0.1% Triton X in PBS) for 45 min, followed by incubation with primary antibody(UCHL1(Invitrogen,1:2000); p-αSyn(CST,1:1000)) at 4°C overnight, and incubation with secondary antibody(anti mouse(Invitrogen,1:500); anti rabbit(Invitrogen,1:400)) was done for 1.5 hours. For nuclear staining, DAPI was used for 8 min. After fixation with prolonged gold, slides were visualised using confocal microscopy with a 63X oil immersion objective. For further analysis, Leica TCS SP8in built software, LAS X, and ImageJ software were used.

### Quantification of estradiol in brain tissue

Estradiol extraction was performed using 50 mg of brain tissue, which was homogenised using a hand homogeniser (in 500 ml of PBS), and further cell lysis was performed with probe sonication (50 Amp, 10 pulses, 0.5 cycle). Lysates were spiked with 20 μl of 7 ng/ml of the deuterated form of estradiol (d3E) as an internal standard. Lysate was transferred to a glass tube, and estradiol was extracted using 2 mL of n-butyl chloride. The upper organic layer was transferred to another tube and dried using N_2_ fumes. Estradiol was derivatised using dansyl chloride for better ionisation. Briefly, 70 μl of a 1:1 solution of dansyl chloride(1 mg/ml) and NaHCO_3_ (pH 11.0) was added and heated for 5 min at 65 °C with shaking at 600 rpm. The mixture was centrifuged at 14000 rpm for 10 min, and the supernatant was used for LC-MS analysis.

The data acquisition was performed using a QTRAP6500+ coupled with a UPLC system (Exion LC), in a scheduled MRM scan mode with positive ionisation polarity.

The instrument was operated with a Turbo Spray Ion Drive at an ion spray voltage of 4500 V, source temperature of 300 °C, curtain gas at 20 psi, ion source gas 1 at 20 psi, and ion source gas 2 at 20 psi. Compound-specific parameters included a declustering potential of 70 V, and entrance and exit potentials of 10 V and 15 V, respectively. MRM duration window was kept 100 s with precursor 506.3, product 171.1, retention time 12.12 min, and CE 34.315 V for estradiol and precursor 509.3, product 171.1, retention time 12.05min, and CE 34.465 V for d3E. We used a solvent gradient of 40-98% solvent B (Acetonitrile) over a period of 18 min. The flow rate was maintained at 0.50 ml/min. Data was processed using MultiQuant, and peak picking was performed manually. Intensities of endogenous β-estradiol were normalised with the intensities of spiked d3E. Average normalised intensities from all 3 sets were used for calculating the amount of estradiol present in the brain tissue.

### Statistical Analysis

Data on the graphs are presented as mean values, and error bars indicate the standard error of the mean. Data from individual mice were represented as a single point in all the bar graphs, while in XY-plots, only mean values were used for representation. Statistical analysis was performed in Graphpad Prism 10.3.1 version using one-way ANOVA with multiple comparisons by comparing the mean from each column. All the analyses were performed among the groups of the same sex. No comparison was conducted between male and female groups. Differences were considered significant if p<0.05 in behavioural, western blot, and confocal data, while in proteomics data analysis, values with FDR≤0.05 were considered.

## Data availability

The mass spectrometry data generated in this study have been deposited in the ProteomeXchange database under accession code PXD065148.

## Reviewer access details

Project accession: PXD065148; Token: nPyd760DqLEd

## Acknowledgements

This work is funded by the Regional Centre for Biotechnology, Faridabad, India. We appreciate the continuous support from technical assistance at the Animal Experimental Facility (AEF) in maintaining and providing transgenic mice. We thank the RCB mass spectrometry facility and the microscopy facility for data acquisition. SA and NK thank the Council of Scientific and Industrial Research (CSIR), Government of India, for the fellowship, and AB, CD, and SK thank the Department of Biotechnology, Government of India, for fellowships.

## Author contributions

SA and TKM conceived the project and developed the experimental design. SA performed all in vivo experiments. AB and NK provided expertise on proteomics sample acquisition and data analysis. NS provided expertise in the quantification of metabolites using targeted-MS. KSB, CD, and SK supported animal experiments and behavioral studies. SA wrote the manuscript, and TKM and AT edited the manuscript. All authors read and approved the final manuscript.

## Additional information

Supplementary Figure S1 and Figure S2.

