## Supplementary Information for "Synapse-related protein alterations and estradiol deficiency associate with early Parkinsonism in female A53T-α-synuclein transgenic mice fed on a high-fat diet"

**Figure S1**.

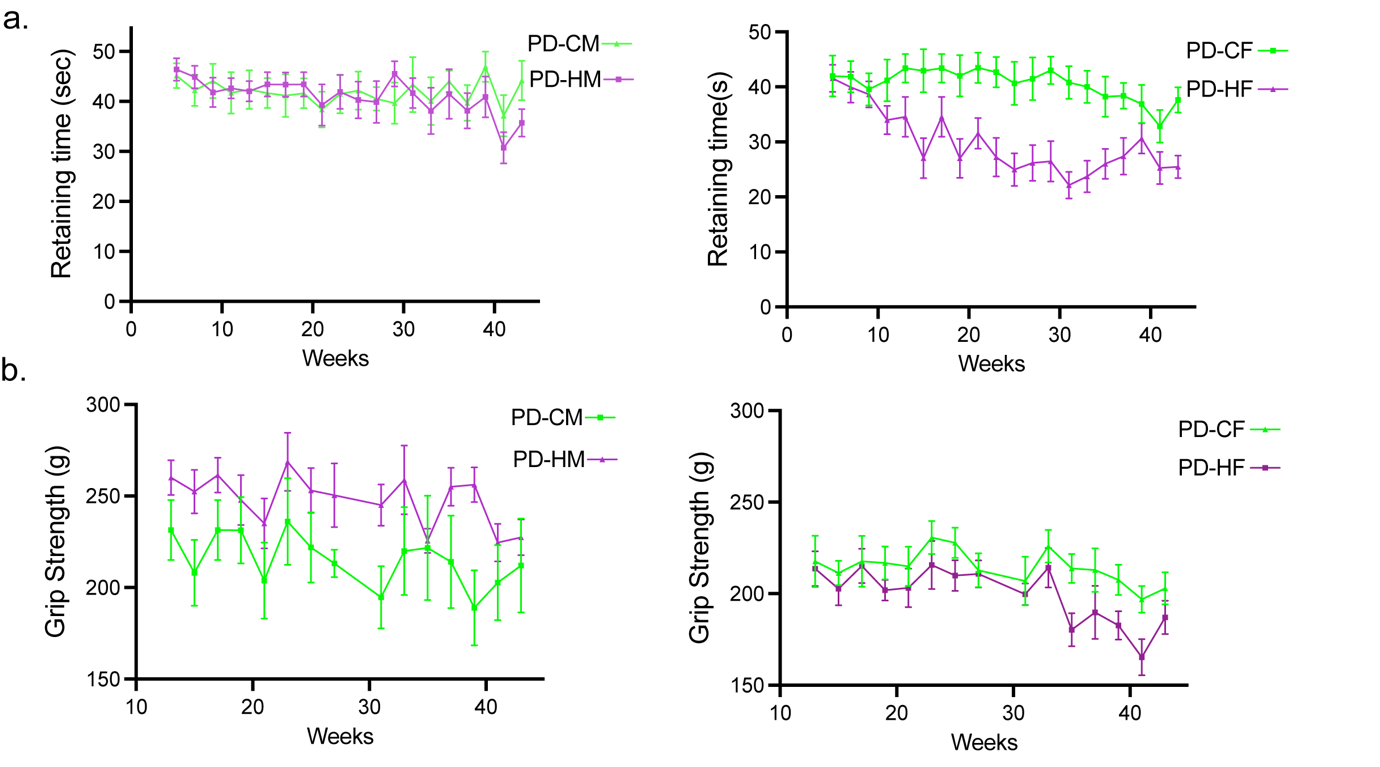

### Fig.S1: Gradual changes in PD progression indicating early PD progression in females under the influence of HFD. a. The graphs show changes in the retention time in the rota-rod measured once every fortnight, starting from the age of 5.5 months till euthanisation in males(left) and females(right) ( n = 7 for PD-CM, n=8 PD-HM, n = 8 for PD-CF, n=9 PD-HF). b. The graphs indicated changes in grip strength measured once every fortnight starting from the age of 5.5 months till euthanisation in males(left) and females(right). ( n = 7 for PD-CM, n=8 PD-HM, n = 8 for PD-CF, n=9 PD-HF). Data are represented as mean ± SEM.

**Figure S2.**

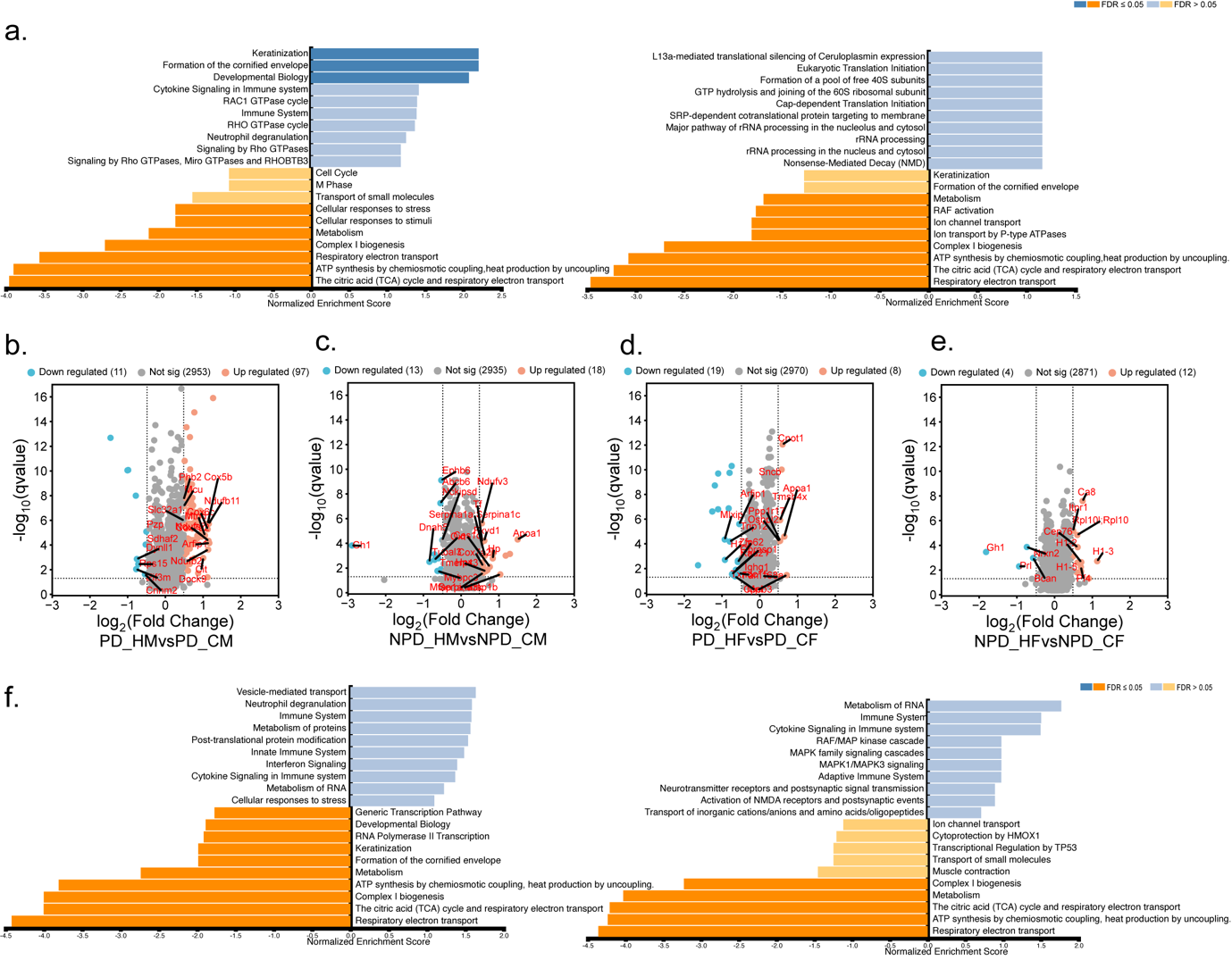

**Fig. S2.** **Independent effect of incorporation of A53T*SNCA and Diet.** . a. Pathways enriched using the fold change of DEPs from the comparison between PD and NPD groups fed on a control diet, indicating 10 positively enriched and 10 negatively enriched pathways in males(left) and in females(right). The X-axis represents the normalised enrichment score of the pathways. b. Volcano plot for the comparison between male PD mice fed on a control diet and HFD, indicating total identified protein along with differentially regulated proteins(DEPs) , x-axis represents log2 fold change, and Y-axis represents. c. Volcano plot for the comparison between male NPD mice fed on a control diet and HFD. d. Volcano plot for the comparison between female PD mice fed on a control diet and HFD e. Volcano plot for the comparison between female NPD mice fed on a control diet and HFD f. Pathways enriched using the fold change of DEPs from the comparison between PD and NPD groups fed on a HFD, indicating 10 positively enriched and 10 negatively enriched pathways in males(left) and in females(right). All belongs to FDR> 0.05
